# Facile synthesis of crosslinked Cu:ZnS-lignocellulose nanocomposite: a potent antifungal and antisporulant system against the tea pathogen *Exobasidium vexans*

**DOI:** 10.1101/2021.09.19.460944

**Authors:** Chayanika Chaliha, Julie Baruah, Eeshan Kalita

## Abstract

The objective of the present study was to synthesize Cu doped ZnS nanocore crosslinked with lignocellulose (represented as Cu:ZnS-lignocellulose nanocomposite) for antifungal action against the devastating tea blister blight pathogen *Exobasidium vexans*. The characteristic features of the nanocomposite were analyzed via different physicochemical techniques like FTIR, XRD, XPS, SEM, SEM-EDX, Elemental mapping, PCS, and UV-PL studies. The FTIR and XPS investigations revealed the crosslinking between lignocellulose and the Cu:ZnS. The presence of lignocellulose was seen to attribute a potent antifungal efficacy, also enhancing the stability of the resulting nanocomposite in aqueous suspensions. The antifungal efficacy confirmed through disk diffusion and broth dilution assays have a maximum zone of inhibition of 1.75 cm^2^ and a MIC50 of 0.05 mg/ml against *E. vexans*. Additionally, the antisporulant activity was evident as the basidiospores failed to germinate in presence of the Cu:ZnS-lignocellulose nanocomposites. This shows potential for stemming the rapid infectivity of *E. vexans* by achieving disease inhibition at the early stage. Finally, the comparison with two commonly used commercial fungicides (copper oxychloride and fluconazole) demonstrated >10-fold higher antifungal activity for Cu:ZnS-lignocellulose nanocomposites.

## 1. Introduction

Nanomaterials have attracted a great deal of attention from researchers for antimicrobial applications owing to their efficient penetration of the microbial cell membrane, targeted delivery, and minimal environmental effects **[1]**. Since the first application of Bordeaux mixture in 1885, Cu ions and their derivatives have long been utilized as defense agents against bacterial and fungal infection in a variety of plant diseases **[2,3]**. Subsequently, numerous nanoparticles have been investigated in the field of plant disease management, with copper-based nanoparticles/nanocomposite being touted as the most potent fungicide. However, the persistent and repetitive usage has raised long-term sustainability concerns that include the emergence of copper-resistant microbial strains; accumulation of Cu in soil, leading to reduced absorption of essential micro/macronutrients by plants; exposure of target crops and non-target soil microbiome to high Cu toxicity, and the retention of copper residues in processed crops posing serious health hazards to consumers **[4]**. Following this, successive legislation introduced by the European Union (EU) now curtail the use of Cu compounds to 4 kg/ha/year **[5]**. Under the new regime, copper formulations such as Cu hydroxides, Cu oxides, and Cu oxychlorides with lower solubility have been approved for usage under strict guidelines, till December 31, 2025 (https://op.europa.eu/s/pzfU). Such low water solubility Cu-based fungicides are prone to the formation of the water-stable film after foliar applications, lowering Cu bioavailability and thus antifungal activity **[6]**. Thereby, the concept of developing copper-based nanoparticles/nanocomposites evolved wherein, nano-sized Cu particles possess enhanced bioavailability, solubility and high surface area to volume ratio and addresses the probable health concerns **[7]**. Besides, there has been some progress in generating Zn-based nanomaterials which have shown promise in mitigating phytopathogenic growth, while being environmentally and economically sustainable **[8, 9]**.

The recent trend of engineered nanomaterials, show considerable success with the usage of doped nanosystems wherein the Cu or Zn entities are doped onto other nanosystems such that they can be effective while being used in trace quantities **[10-12]**. However, the potential of such doped nanosystems against the robust phytopathogenic fungi remains to be fully explored. Lignocellulose biopolymers are now recognized as the futuristic precursors for the generation of sustainable antimicrobial matrices, which can be potent and stable for long durations via stabilizing the nanocore from agglomeration and oxidation **[13, 14]**. These materials have the added advantages of being economical, regenerative, environmentally benign, and biodegradable **[15]**. Taking into account the aforementioned considerations, the scope of the current study is to develop a multimodal antifungal system *viz*. Cu doped ZnS (Cu: ZnS) nanocomposite crosslinked within a lignocellulose matrix. The synthesized antifungal agent aspires to deliver the potent efficacy of conventional Cu-based antimicrobial against the blister blight causal organism of tea *Exobasidium vexans*, using trace amounts of Cu in a chemically stable formulation. For synthesis, a benign surfactant, sodium citrate, was used to prevent agglomeration of the Cu: ZnS nanomaterials prior to its crosslinking within the lignocellulose matrix. The Cu: ZnS-lignocellulose nanocomposites has been subsequently evaluated for their antifungal and antisporulant efficacies against the recalcitrant phytopathogen *E. vexans*.

## 2. Materials and Methods

### 2.1. Chemicals

Analytical grade reagents *viz*. Copper acetate (Cu (CH_3_COO)_2_), Zinc chloride (ZnCl_2_), Sodium sulfide (Na_2_S), Sodium borohydride (NaBH_4_), and Sodium citrate (NaC_6_H_5_O_7_) were procured from Sigma-Aldrich India. Rice husk samples were employed as the precursor for lignocellulose and were collected from local rice mills of Tezpur, Assam, India.

### 2.2. Synthesis of lignocellulose based Cu:ZnS nanoparticles

Synthesis of Cu:ZnS-lignocellulose nanocomposites were carried out based on our earlier work, with appropriate modifications (Fig. 1 a) **[16]**. Stoichiometric amounts of ZnCl_2_ (16mM), Na_2_S (50 mM), and Cu (CH_3_COO)_2_ (4mM), and were dissolved in distilled water to obtain the solvothermal premix followed by the addition of Sodium citrate (19mM). Rice husk was washed vigorously to remove surface impurities, oven-dried (50°C), and pulverized at a frequency of 8.33 Hz in a planetary ball mill. The milled rice husk (2% w/v) was then added to the premix, followed by the gradual addition of NaBH_4_ (75mM) under constant stirring. The solvothermal treatment of the entire reaction mix was initiated in a round bottom flask at 100°C, to produce time-based variations based on solvothermal residence time (Fig. 1 a). Upon completion, the solvothermal extract was cooled to room temperature, washed several times (water and ethanol), and finally oven-dried (60 °C) for a period of 48 h. The schematic illustration of the synthesis of the Cu:ZnS nanoparticles followed by their crosslinking into the lignocellulose nanocomposites is depicted in Fig. 1 b.

**Fig.1.**
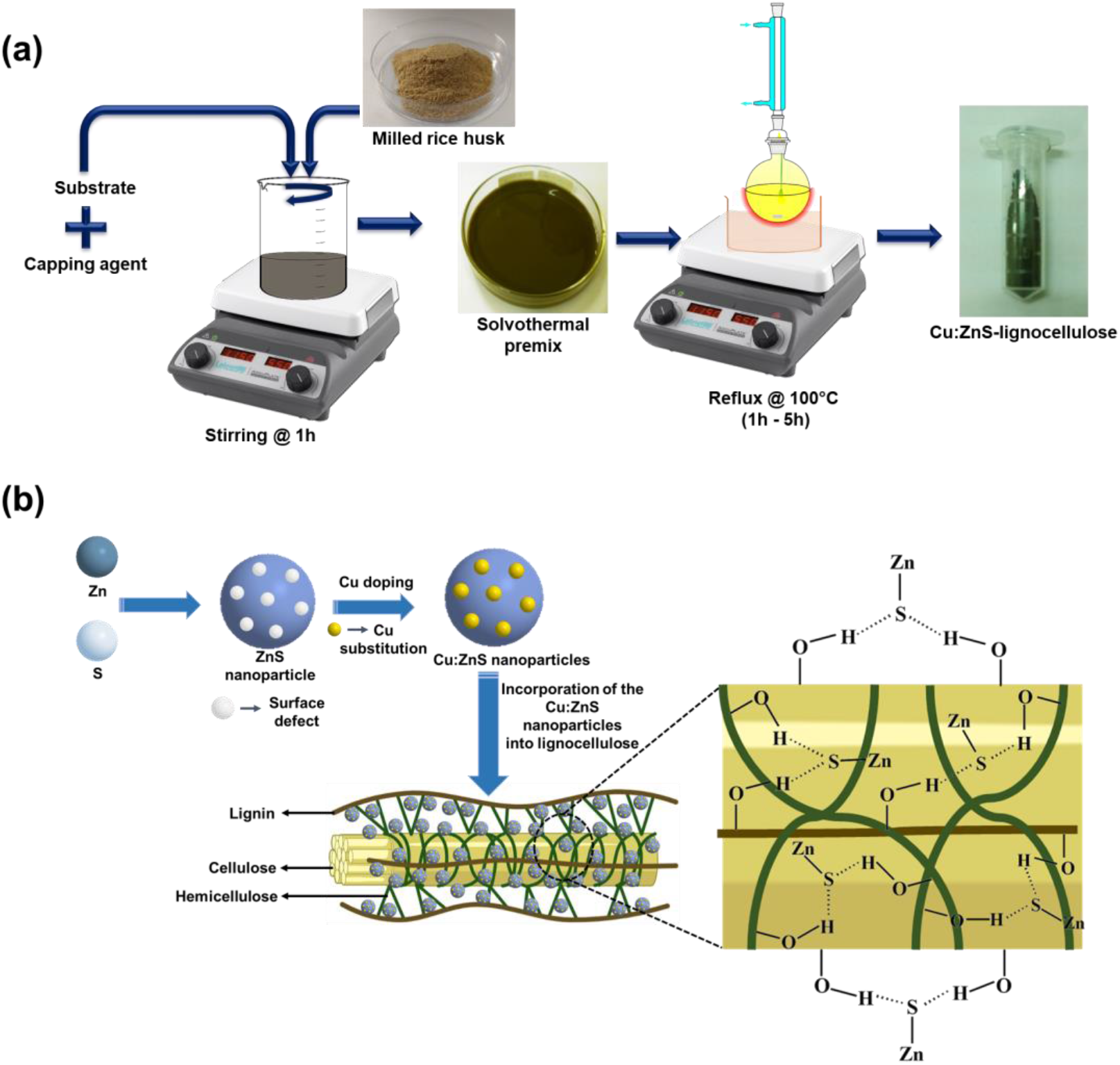
Schematic representation of the synthesis procedure (a) and the formation (b) of Cu:ZnS- lignocellulose nanocomposites. (Created with BioRender.com)

### 2.3. Characterization of synthesized nanocomposites

Fourier-transform Infrared (FTIR) spectra of the nanocomposites were documented on FT-IR Spectrometer (Perkin Elmer-Spectrum 100 Optica, USA) to study the functional group present in the system. The crystalline structures of the synthesized nanocomposites were analyzed on the RIGAKU-Miniflex Benchtop, Japan, X-ray powder diffractometer. To detect the binding energies, X-ray photoelectron spectroscopic analysis was carried out with XPS, Thermo Fisher Scientific Pvt. Ltd., ESCALAB Xi+, UK. The morphology of the synthesized nanocomposites was examined through the Scanning Electron Microscope (SEM) (JSM-6390, JEOL, Singapore) and the elemental composition were analyzed using SEM-EDX (SEM-Energy Dispersive X-ray Diffraction spectroscopy). Pseudo-colors were used to generate the Elemental map for the nanocomposites to illustrate the elemental distribution in the synthesized nanocomposites. Additionally, the absorption spectra of the nanocomposites showing the highest antifungal efficacy were attained in the range of 200–800 nm with a UV-Vis spectrophotometer (Thermo Scientific UV-10). The photoluminescence (PL) excitation spectra of the sample were also noted by a fluorescence spectrophotometer (Perkin Elmer LS55).

### 2.4. Effect of solvent pH on the hydrodynamic diameter and electrostatic stability of the nanocomposites

Since the nanocomposites have been synthesized for foliar application, they were analyzed for their variation in the size in an aqueous dispersion by observing the alterations to their hydrodynamic diameter and electrostatic stability concerning the change in solvent pH. The synthesized nanovariants, dispersed in deionized water with different pH were ultrasonically treated for 10 min at a frequency of 24 kHz (Sartorius labsonic M, Germany). The difference in the hydrodynamic diameter of the nanocomposites was investigated using a Zetasizer Nano-ZS-90 Dynamic Light Scattering (DLS) system (Malvern Instruments, UK) at a wavelength of 633 nm and a scattering angle of 90°, under a constant temperature of 25°C. Aqueous dispersions of the nanocomposites were evaluated for their electrostatic stability by examining their surface charge (ζ potential) with respect to the change in solvent pH using a Zetasizer Nano-ZS-90 DLS system (Malvern Instruments, UK).

### 2.5. Antifungal assay

The synthesized nanocomposites were tested for antifungal properties against *E. vexans* by performing disk diffusion assays. The spore suspension was prepared with a spore count of ∼10^6^spores ml^-1^ from 5-7 days old fungal culture grown on czapek dox media **[17]** and then spread on czapek dox agar (CDA) plates. Sterile filter paper disks were dipped in different concentrations (0.5 mg/ml, 0.8 mg/ml, and 1 mg/ml) of the synthesized nanovariants, allowed to dry, and were then placed in the CDA plates. The plates were incubated for 5 days at 28°C and then examined for the presence of a prominent zone of inhibition. The antifungal activity of the synthesized nanocomposites was compared with the conventional commercial fungicides copper oxychloride and fluconazole. For estimating the minimum inhibitory concentration (MIC) and minimum fungicidal concentration (MFC) values, different concentration of the antifungal agents (0.05 mg/ml, 0.08 mg/ml, 0.1 mg/ml, 0.25 mg/ml and 0.5 mg/ml) were added to a 96-well microtitration plate containing czapek dox broth media. The wells were subsequently inoculated with a spore suspension of ∼10^6^ spores ml^-1^ and incubated at 28°C for 48 h. The MIC_50_ was recorded for the lowest antifungal concentration at which 50% growth was observed **[18]**. For MFC calculations, aliquots from each well at the aforementioned concentrations were taken spread on CDA plates, which were incubated at 28°C for 48 h. The MFC value was recorded for the lowest antifungal concentration where less than three fungal colonies were observed.

### 2.6. Antisporulant activity

To determine the antisporulant activity, the inhibitory effects of the synthesized nanocomposites on basidiospore germination of *E. vexans* were evaluated by the concavity slide method **[19]**. Briefly, 100 µl of basidiospore suspension (∼10^6^ spores µl^-1^) was suspended with 2 h variant of the Cu:ZnS-lignocellulose nanocomposite suspension at different concentrations (0.5 mg/ml, 0.8 mg/ml, and 1 mg/ml). 50 µl of the mixed suspensions were placed in slides, incubated for 8 h at 28°C, and were observed for germination and hyphal growth under a microscope (Olympus BX 43). Basidiospore suspensions placed on slides without the addition of nanocomposites were considered as a blank. For scoring, the germ tube extension to twice the length of basidiospores was considered as germinated basidiospores. The rate of inhibition of basidiospore germination was determined as the ratio between the number of non-germinating basidiospores and the sum of spores observed under a light microscope for five independent microscopic fields of vision.

## 3. Results and discussion

### 3.1. Characterization of synthesized nanocomposites

The structural and crystalline attributes of the Cu:ZnS-lignocellulose nanocomposites (1 h-5 h variants) and the native Cu:ZnS nanocomposite (2 h variant), were analyzed through FTIR and XRD techniques (Fig. 2 a, b). In Fig 2a, demonstrates the dominant absorption band at 545 cm^-1^ and 976 cm^-1^ corresponding to the ZnS vibration and resonance interaction between Cu and Zn respectively **[16]**. Additionally the prominent peak at 1218 cm^-1^ for the Cu:ZnS-lignocellulose nanovariants corresponds to CH_2_ vibrations at C-6 or O-C-stretching of cellulose I suggesting an interaction between lignocellulose and ZnS **[20, 21]**. The crosslinking between the -OH group of lignocellulose and ZnS through hydrogen bonding, is evident from the shifting of the characteristic band for the inter and intramolecular -OH stretching vibration (3401 cm^-1^) for the Cu:ZnS nanocomposite to 3428 cm^-1^ in Fig. 1 b **[22]**. The band observed at 1450 cm^-1^ corresponds to the aromatic skeletal vibration of the phenolic group of lignin **[23, 24]**. Also, the characteristic signature for the presence of capping agent SC is represented by the band at 1601 cm^-1^ which corresponds to the stretching vibration of the citrate group **[25]**. Fig. 2 b depicts the XRD diffractograms of the Cu:ZnS-lignocellulose nanovariants (1 h – 5 h) and 2 h variant of Cu:ZnS nanocomposite for all the reflux variants. The diffractograms for Cu:ZnS and Cu:ZnS-lignocellulose nanocomposites display characteristic peaks at 28°, 47°, and 58° corresponding to (111), (220), and (311) diffraction plane respectively which are in good agreement with the cubic Zinc blende phase structure of ZnS for the synthesized variants **[26]**. However, an additional peak at 34.5° corresponding to the (004) diffraction planes respectively was found in Cu:ZnS-lignocellulose nanovariants indicating major characteristic diffraction peaks of typical cellulose I structure **[27]**. The lower intensity of peaks in the Cu:ZnS-lignocellulose nanovariants compared to Cu:ZnS signifies the reduction in crystallinity, owing to the crosslinking with the amorphous lignocellulose matrix **[28, 29]**.

**Fig. 2.**
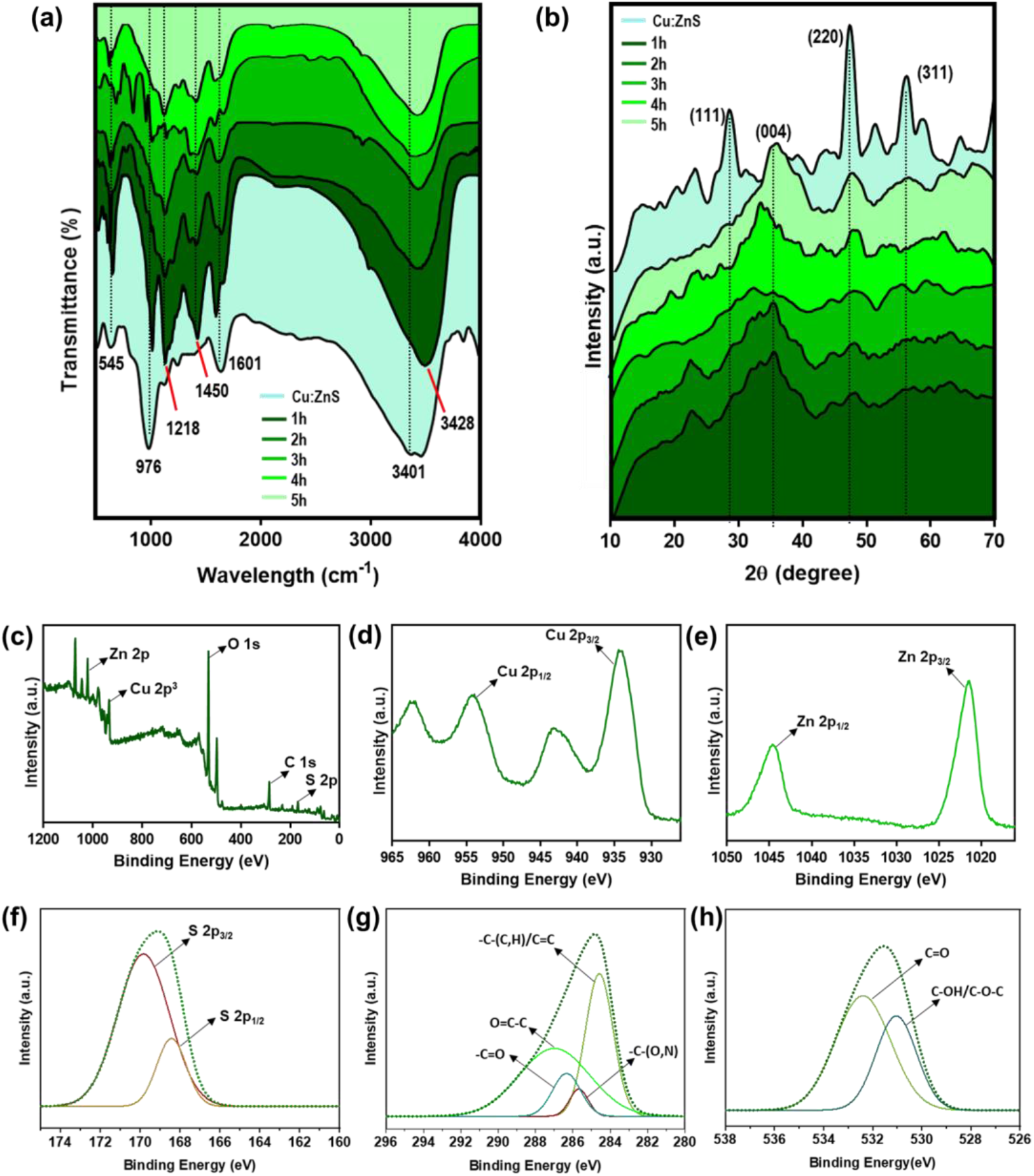
FTIR spectra (a) and X-ray diffraction spectra (b) of all the variants of Cu:ZnS- lignocellulose nanocomposites (1 h – 5 h) and 2 h variant of Cu:ZnS. XPS spectrum of 2 h variant (c), high-resolution XPS scan for Cu 2p_3_ (d), Zn 2p (e), S 2p (f) C 1s (g), and O 1s (h) of 2 h variant of Cu:ZnS-lignocellulose nanocomposite.

To investigate the chemical states of the elements and to characterize the functional groups present in lignocellulose XPS analyses were performed and the XPS results are depicted in Fig. 2 c. The XPS survey spectra of the 2 h variant of Cu:ZnS-lignocellulose nanocomposite show the peaks of the elements Zn, Cu, S, C, and O. The spectrum shows the Cu 2p^3^ binding energy values at 934.03 eV for the presence of copper. The peak at binding energy 1021.08 eV and 168.93 eV corresponds to the Zn 2p and S 2p in the sulfide phase respectively, corroborating the existence of ZnS (Fig. 2 c) **[22]**. The constituent elements C and O for the lignocellulose component of the nanocomposite were confirmed with the presence of photoelectron peaks at binding energy 540.5 and 293.5 eV corresponding to O 1s and C 1s respectively (Fig. 2 c). The elemental specific spectra of Cu have been depicted in Fig. 2 d. The photoelectron peak at binding energy 934.24 eV and 954.20 eV of Cu spectra indicates the formation of Cu^2+^ oxidation state. However, the additional peak around 940 eV and 960 eV results from a shakeup process owing to the open 3d^9^ shell of Cu^2+^ [22]. The elemental spectra for Zn show peaks at 1021.43 and 1044.65 eV corresponding to Zn 2p^3/2^ and Zn 2p^½^ belonging to the Zn^2+^ oxidation state (Fig. 2 e) **[22]**. The binding energies 168.4 eV and 169.6 eV) in Fig. 2 f correspond to S 2p_3/2_ and S 2p_1/2_, in accordance with the binding energies for sulfur in metal sulfides (M-S), indicating the formation of ZnS in the nanocomposite **[30]**. The high-resolution C 1s spectra were resolved into four individual component peaks at 284.3 eV, 285.8 eV, 286.2 eV, and 287 eV corresponding to –C-(C, H)/C=C (i.e. C singly-bound to C/H or doubly bound to C), -C-(O, N) (C bound to C or N), -C=O (C doubly bond to O) and O-C=O (C singly and doubly bound to O), respectively (Fig. 2 g) **[31]**. In contrast, the O 1s XPS spectra exhibit two deconvoluted peaks at 533.1 and 531.7 eV, signifying C=O and C–OH/C–O–C groups, respectively (Fig. 2 h) **[32]**. The existence of these functional groups in the O 1s XPS spectra is consistent with the FTIR results, indicating the presence of O-C-stretching vibrations supporting the crosslinking reaction scheme between lignocellulose and ZnS **[21]**. The elemental composition of the 2 h variant of Cu:ZnS-lignocellulose nanocomposite obtained from XPS analysis are listed in Table S1.

The morphological structure of the surface of the 2 h variant of the synthesized nanocomposite was obtained with SEM-EDX analysis. The SEM micrograph for the 2 h variant of Cu:ZnS-lignocellulose nanocomposite (Fig. 3 a) shows a quasi-spherical morphology. The existence of C, O, S, Cu, and Zn as major elemental constituents in the nanocomposite was confirmed from SEM-EDX studies (Fig. 3 b) and the weight % and atomic % of the elemental compositions are listed in Fig. 3 b inset. Pseudo-color elemental mapping for the 2 h variant was carried out wherein C, O, S, Cu, and Zn demonstrate a uniform distribution of the elements within the area under investigation (Fig. S1 a-e).

**Fig. 3.**
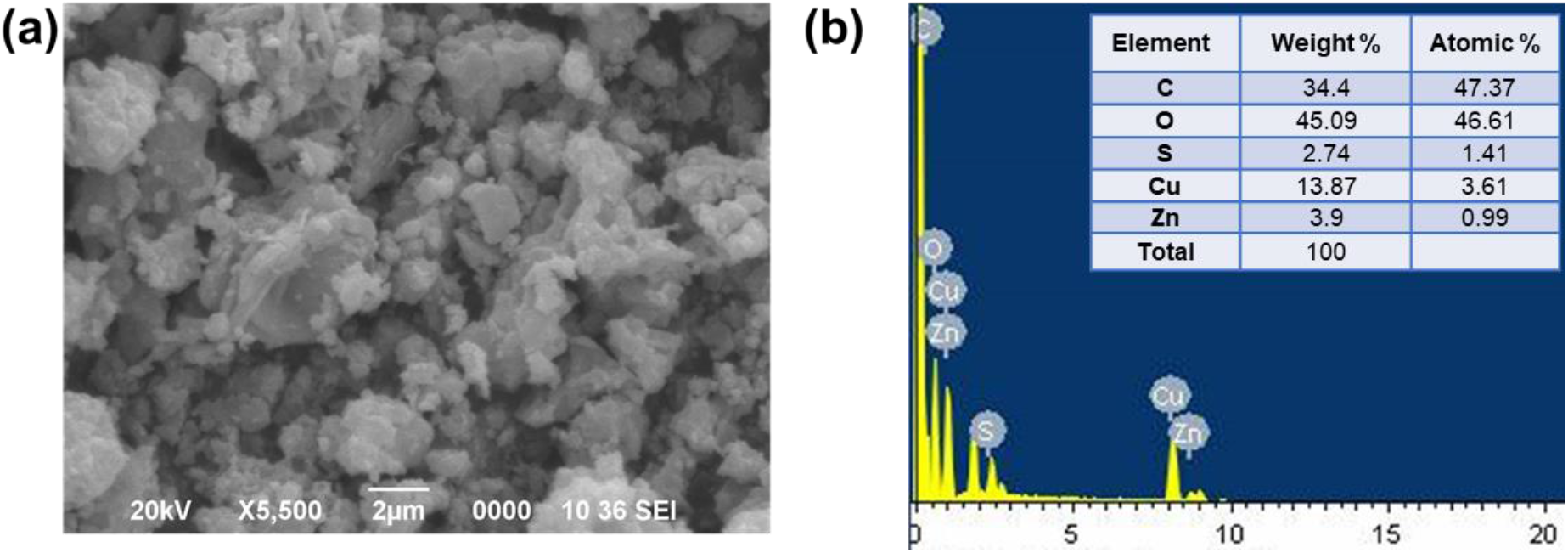
SEM micrographs (a) and EDX spectra (b) with elemental composition (Inset b) of 2 h variant of Cu:ZnS-lignocellulose nanocomposite.

### 3.2. Effect of solvent pH on the hydrodynamic diameter and electrostatic stability of the synthesized nanocomposites

The particle size distributions and electrostatic stability of Cu:ZnS-lignocellulose nanocomposites under different solvent pH, were estimated using Photon Correlation Spectroscopy (PCS) to analyze the particle behaviour (Fig. 4 a, b). From Fig 4a. it is evident that there is no significant change in the hydrodynamic diameter of the different variants, across the pH range indicating their stability across pH 2-12. It has been reported that colloidal dispersions with a ζ potential of ≥ ±20 meV are generally regarded stable **[33]**. The ζ potential of the synthesized nanocomposites were found to be in the -22 meV to -34 meV range (Fig. 4 b), across the solvent pH 2-12. Here, the 2 h reflux variant was found to be the most stable with the highest ζ potential. Further, the sizes of the variants were well within the desirable range and is indicative of their well dispersed state in the solvent. The high electrostatic stability and stable hydrodynamic diameter of the dispersed nanocomposite could be attributed to the crosslinking of Cu:ZnS core to the lignocellulose matrix. This is likely to address the concerns of low water solubility, oxidation, and agglomeration associated with the conventional copper-based fungicidal formulations.

**Fig. 4.**
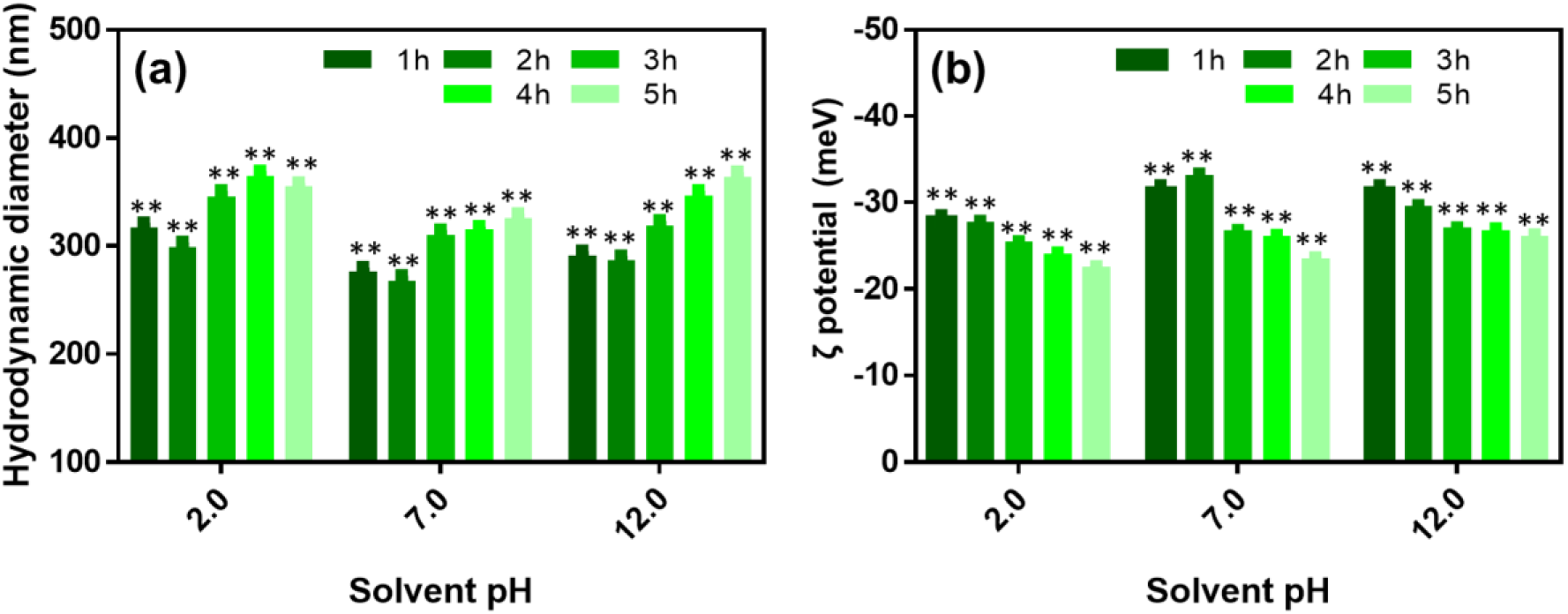
Hydrodynamic diameter and ζ potential of Cu:ZnS-lignocellulose nanocomposites across solvent pH [**P-value 0.001-0.01].

### 3.3. Antifungal assay

The as-synthesized Cu:ZnS-lignocellulose nanocomposites were initially evaluated for their antifungal efficacy by performing the disk diffusion assay. It is evident from the disk diffusion assays, that the nanocomposites when used at concentrations of 0.5 mg/ml, 0.8 mg/ml and 1 mg/ml, result in a significant concentration dependent inhibition of *E. vexans* hyphal growth (Fig. 5 a). Among the nanocomposites, the 2 h variant of Cu:ZnS-lignocellulose shows the (Fig. 5 b) maximal antifungal efficacy with the zone of inhibition measuring ∼1.76 cm^2^. In comparative studies with the common commercial fungicides for blister blight control, *viz*. copper oxychloride and fluconazole, Cu:ZnS-lignocellulose nanocomposites also show better antifungal efficacy **[34]**. The nanocomposites demonstrate the zone of inhibition for concentrations as low as 0.5 mg/ml, while copper oxychloride is effective only at a concentration of 4 mg/ml (Fig. 5 b). On the other hand, fluconazole was unable to inhibit *E. vexans* growth even at the concentration of 4 mg/ml. Further, the MIC_50_ and MFC (48 h) values derived for the nanocomposite variants are depicted in Fig. 5 c. The lowest MIC_50_ of Cu:ZnS-lignocellulose nanocomposites, which inhibited *E. vexans* growth by 50%, was achieved at concentration 0.05 mg/ml for the 2 h variant. Likewise, the lowest MFC value was at the concentration 0.25 mg/ml for the 2 h variant of Cu:ZnS-lignocellulose. In addition to this an activity index of the nanocomposite variants was calculated at a concentration of 4 mg/ml, based on the antifungal activity of copper oxychloride (Fig. 5 d). A >10-fold increase in the activity was observed for the 1 h - 4 h variants of the nanocomposites (Fig. 5 d). Here the 2 h variant of Cu:ZnS-lignocellulose, was most potent as it exhibited a **>**12 fold increase in activity against *E. vexans*.

**Fig. 5.**
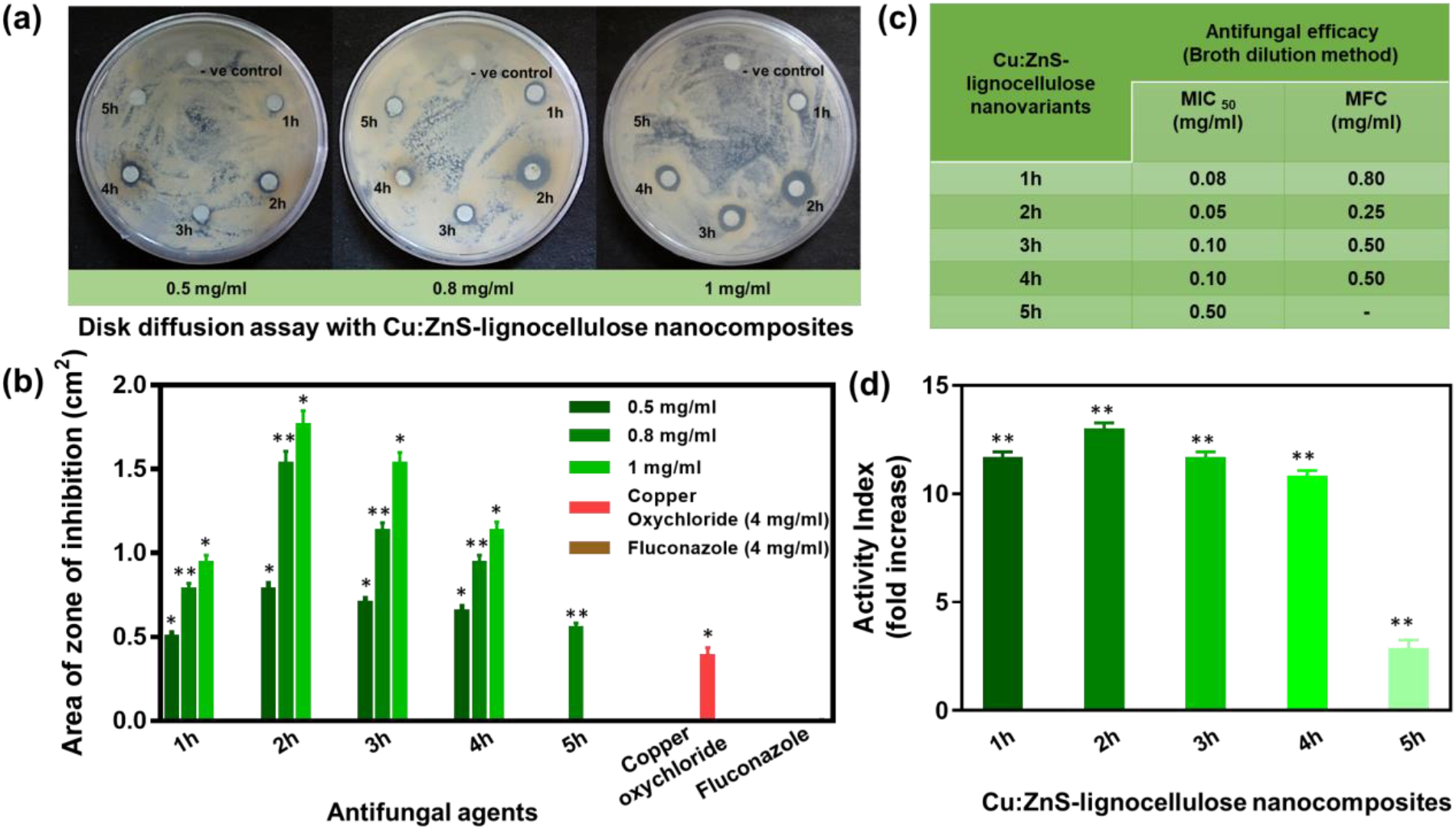
Disk diffusion assay showing antifungal efficacy of Cu:ZnS-lignocellulose nanocomposites (a), graphical representation of antifungal efficacy of Cu:ZnS-lignocellulose nanocomposites, copper oxychloride and fluconazole (b), MIC_50_ and 48 h MFC values (c) and activity index (d) for Cu:ZnS-lignocellulose nanocomposites against *E. vexans* [**P-value 0.001-0.01, *P-value 0.01- 0.05].

### 3.4. Antisporulant activity

The basidiospores, as a propagative unit, considerably aid in the rapid proliferation and pathogenicity of *E. vexans* on host tea. Basidiospore germination being the first significant phase in blister blight infestation, its suppression is linked to inhibition of disease at the early stages. To further appraise the fungicidal efficacy, the 2 h Cu:ZnS-lignocellulose nanocomposite variant, having the highest activity index was investigated for its antisporulant activity. Fig. 6 (a-d) shows the light micrographs depicting the germination of basidiospore after incubation with the 2h Cu:ZnS-lignocellulose nanocomposite suspensions and under control conditions. As expected, the control samples (basidiospores in sterile water) showed the germination of basidiospores with extensive hyphal growth 8 h post-incubation (Fig. 6 a). The presence of the Cu:ZnS-lignocellulose nanocomposite resulted in arrested germination and hyphal growth of *E. vexans* basidiospores in a concentration dependent manner (Fig. 6 b-d). As such, the highest concentration of 1 mg/ml, completely inhibited the *E. vexans* basidiospore germination, while also exhibiting sporicidal effects. Fig. 6 e depicts the calculated rate of inhibition of basidiospore germination when incubated with and without nanocomposites. An inhibition of 0%, 65%, 80% and 100% is evident for the concentrations 0, 0.5, 0.8 and 1mg/ml, respectively. To determine the efficacy of antisporulant activity a scale was introduced, based on the microscopic observations, to grade the activity wherein -, + and ++ indicated no inhibition, partial inhibition, and full inhibition respectively. The score of inhibition for the 2 h variant of Cu:ZnS-lignocellulose nanocomposite at different concentrations against basidiospore of *E. vexans* is shown in Fig. 6 f.

**Fig. 6.**
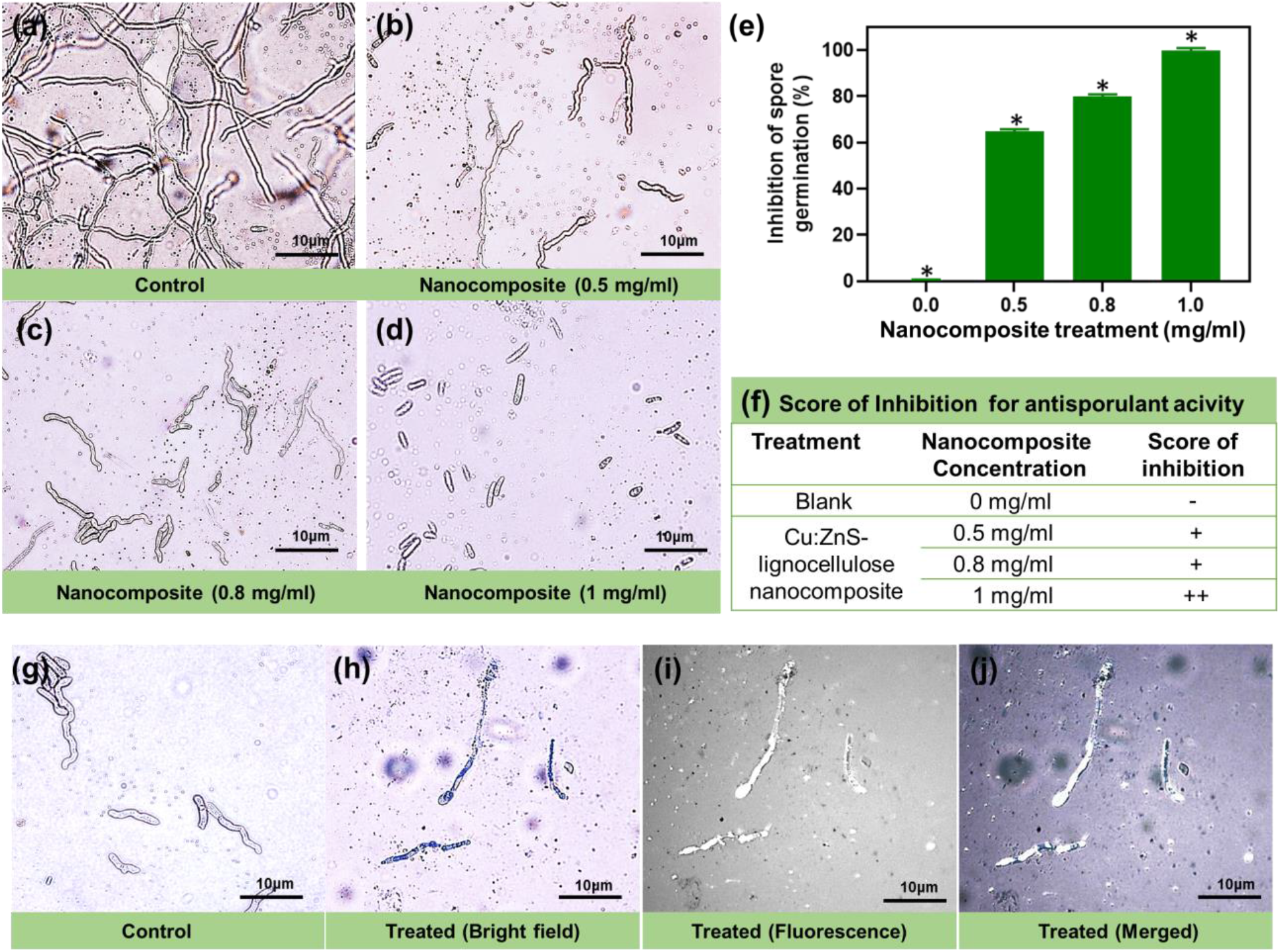
Antisporulant activity. Germination of *E. vexans* basidiospore incubated in water (Control) and 2 h variant of Cu:ZnS-lignocellulose nanocomposite suspension (a-d), Rate of inhibition of basidiospore germination at different concentration of 2 h variant of Cu:ZnS-lignocellulose nanocomposite suspension [*P-value: 0.01-0.05] (e), Scoring of antisporulant acitivity [Scale: no inhibition (-), partial inhibition (+) & full inhibition (++)] (f), untreated basidiospore morphology observed under bright field microscopy (Control) (g), impaired surface of germinating basidiospore observed under bright field microscopy (h), nanocomposites deposition on basidiospore surface observed under fluorescent microscopy (i) and merged image of bright field and fluorescent microscopy (j).

The 2 h variant of Cu:ZnS-lignocellulose nanocomposite was evaluated for its optical/fluorescent properties using UV-PL analysis. The absorption peak was obtained at a wavelength of 205 nm, while the emission peak appeared at the excitation wavelength of 390 and 425 nm (Fig. S2 a, b). The photoluminescence (PL) excitation spectra of the nanocomposite corroborated the fluorescent property of the variant and provided the avenue for the fluorescent microscopic studies of the nanocomposite. To investigate the effect of the nanocomposites on the *E. vexans* hyphal growth, the morphological alterations were microscopically monitored (light and fluorescent) in the presence of 2 h variant of Cu:ZnS-lignocellulose nanocomposites and control conditions. As can be seen in Fig 6 g the germinated hyphae from the control samples retain a complete and homogeneous tube-like morphology. Nanocomposite treatment of *E. vexans* basidiospores resulted in adverse morphological alterations starting with the deposition of the nanocomposites on the germ tube leading to membrane disruption is evident in the bright field and fluorescent micrographs (Fig. 6 h-i). Fig. 6 j represents the merged image of bright field and fluorescent micrograph wherein we can confirm the membrane impairment of the basidiospores owing to the deposition of the nanocomposites.

With respect to the antifungal effect of the Cu:ZnS-lignocellulose nanocomposites it is worth mentioning that the net negative charge of the nanocomposites facilitates its adherence to positively charged chitin of the fungal cell wall through electrostatic interaction. Following the adherence, the anti-fungal activity is likely mediated by the classical pathway of cell wall disintegration, electrolytic imbalance, inhibition of nucleic acid, and protein synthesis impairment (Fig. 7) **[35, 36]**. Cu-based antifungal agents are widely reported to induce oxidative stress on fungal cell walls leading to the disruption of cellular membrane and leakage of cytoplasmic components. One of the primary causes of membrane disruption is lipid peroxidation which is induced by reactive oxygen species (ROS). This formation of reactive oxygen species (ROS) is mediated by a variety of mechanisms including the generation of superoxide via inhibition of the mitochondrial electron-transport chain and the inactivation of enzymes to generate hydrogen peroxide **[37, 38]**. The presence of ROS in turn is known to cause DNA damage and impairment of proteins involved in DNA replication process, in addition to the disruption of the cellular membrane. The micrographs present strong evidence of the membrane adherence of the Cu:ZnS-lignocellulose nanocomposites followed by the membrane disintegration which can be inferred from the morphological deformation of the *E. vexans* basidiospore and germ tube (Fig. 6 j).

**Fig. 7.**
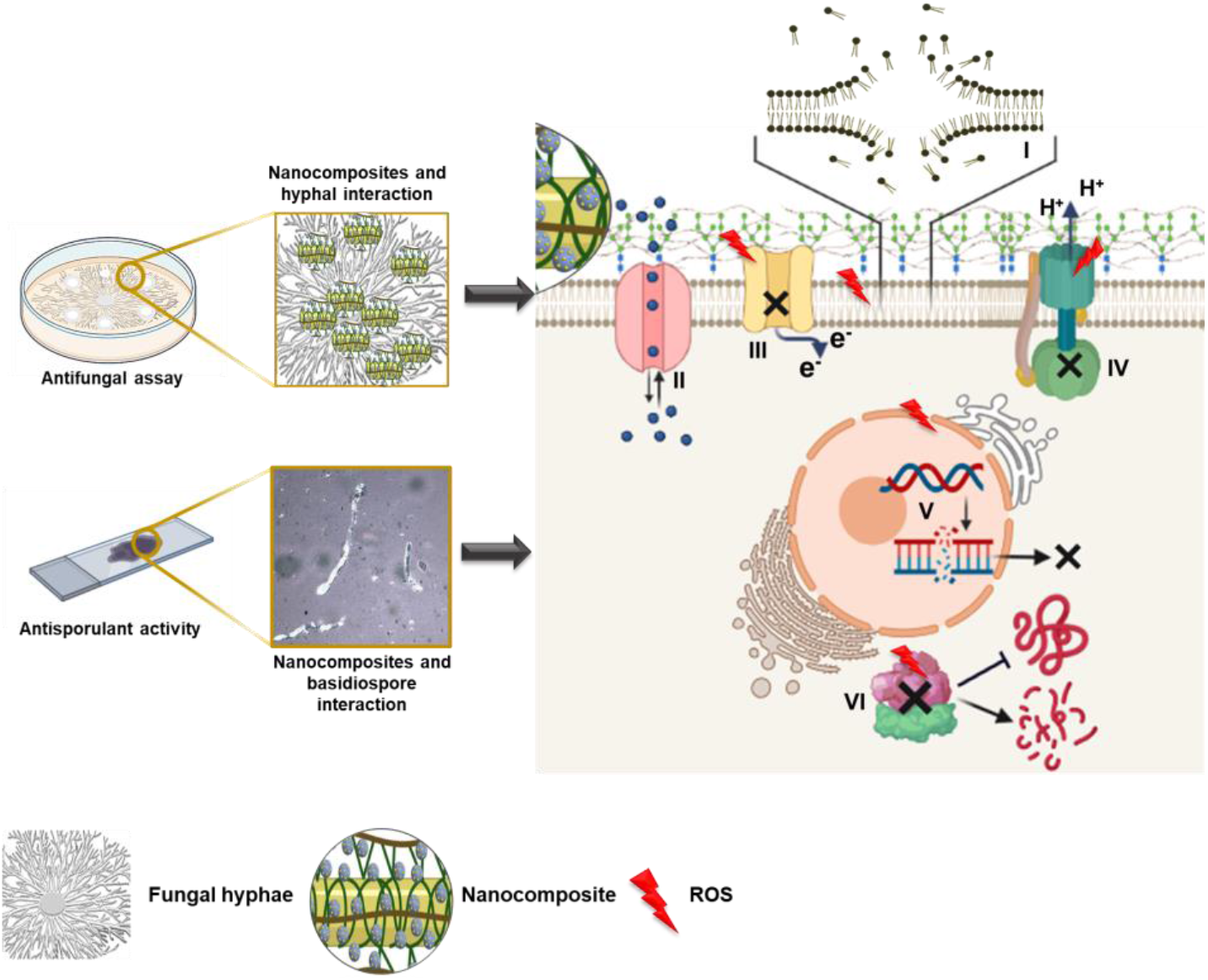
Schematic illustration of Cu:ZnS-lignocellulose nanocomposite triggered antifungal and antisporulant mechanisms: Upon cell surface adhesion of nanocomposites on basidiospores and hyphae, generation of ROS leads to the disintegration of the cell membrane (I), which disrupts membrane permeability (II), electrolytic imbalance (III), and interferes proton pumping (IV). Further, intracellular damages resulting from ROS generation include inhibition of DNA replication (V), and also interferes with ribosomal integration thereby inhibiting protein synthesis and denaturation (VI). (Created with BioRender.com)

## 4. Conclusion

This work demonstrates the antifungal and anti-sporulant potential of the doped Cu:ZnS nanoparticles crosslinked within a lignocellulose to form the Cu:ZnS-lignocellulose nanocomposites, in controlling the blister blight disease. The nanocomposites are established to be more potent than the commercially used fungicides (copper oxychloride and fluconazole) whilst addressing the concerns of bioavailability of copper and copper toxicity. The Cu:ZnS-lignocellulose nanocomposites are electrostatically stable aqueous suspension across a wide pH range making them acceptable for usage as foliar spray. Thus, Cu:ZnS-lignocellulose nanocomposites with its significant antifungal and antisporulant characteristics, can be assigned as a potential candidate for development of an alternative copper-based formulation for the control of the devastating blister blight disease of tea.

## Supporting information

ESM_1

## Acknowledgment

The authors wish to acknowledge DBT, Govt. of India, for the Twinning Research Grant (Grant No. BT/427/NE/TBP/2013). Author CC would like to acknowledge DST, Govt. of India for her DST INSPIRE Junior Research Fellowship (IF-150964). The authors thank Ananda Tea Estate, North Lakhimpur District, Assam, India, for providing the blister blight infected tea leaf samples used in the study.

## Declarations

### Availability of data and material

Not applicable

### Code availability

Not applicable

### Authors’ contributions

**Chayanika Chaliha:**Methodology, Investigation, Formal analysis, Data curation, Writing - original draft. **Julie Baruah:** Formal analysis, Data curation, Writing - review & editing. **Eeshan Kalita:** Conceptualization, Methodology, Supervision, Verification, Writing - review & editing, Project administration, Funding acquisition.

### Author approvals

The authors CC, JB, and EK have seen and approved the manuscript, and it has’nt been accepted and published elsewhere.

