## Supplementary material for "Facile synthesis of crosslinked Cu:ZnS-lignocellulose nanocomposite: a potent antifungal and antisporulant system against the tea pathogen *Exobasidium vexans*": ESM_1

Dr. Eeshan Kalita

Table S1. Elemental composition of 2 h variant from XPS analysis

| **Element** | **Atomic %**  Cu:ZnS-lignocellulose |
| --- | --- |
| **C** | 31.54 |
| **O** | 51.24 |
| **S** | 6.02 |
| **Cu** | 7.19 |
| **Zn** | 4 |

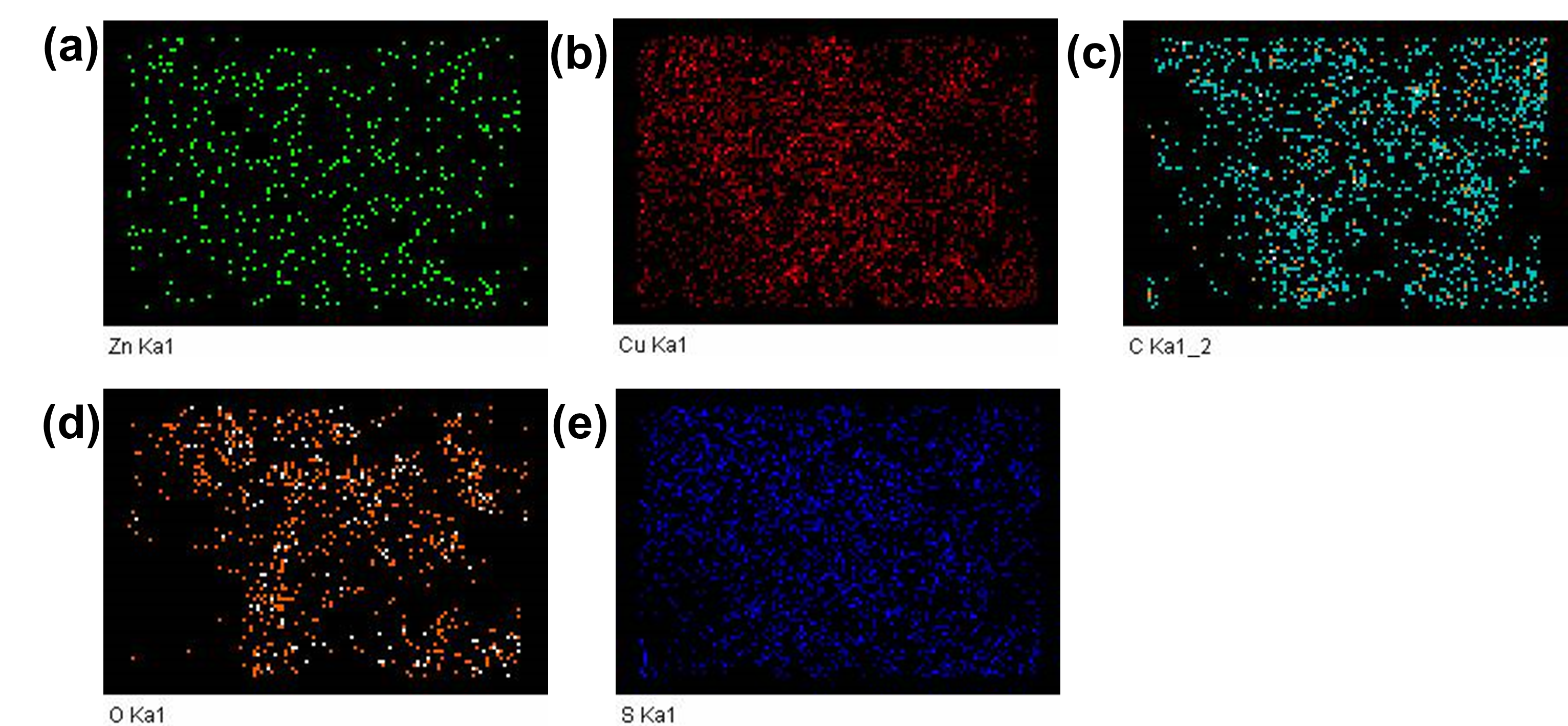

Fig. S1. Pseudo-colour elemental distribution maps of 2 h variant of Cu:ZnS-lignocellulose nanomatrix (a-e).

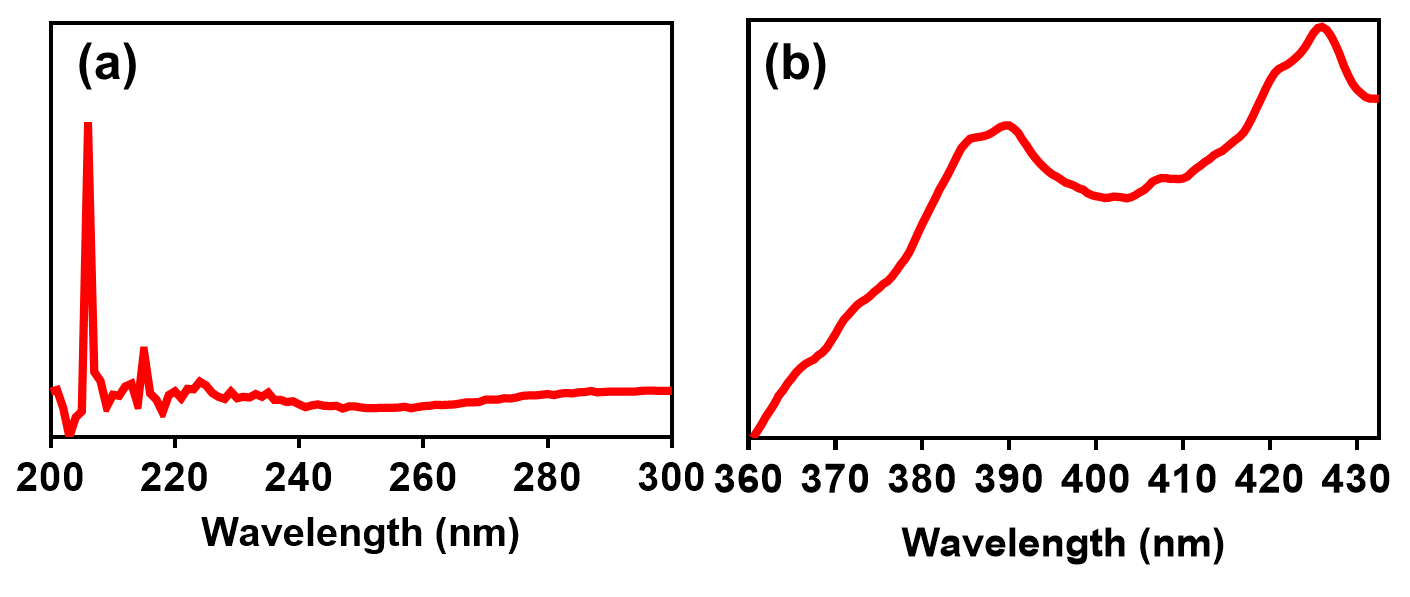

Fig. S2. UV–vis absorption spectra (a) and PL spectra (b) of 2 h variant of Cu:ZnS-lignocellulose nanomatrix
